# Determination of the cellular target engagement by direct-to-biology Cellular Thermal Shift Assay (CETSA)

**DOI:** 10.64898/2026.02.28.708721

**Authors:** Magdalena M. Szewczyk, Darshil Patel, Sarvatit Patel, Aiymzhan Istayeva, Robert Batey, Dalia Barsyte-Lovejoy, Vijayaratnam Santhakumar

## Abstract

The direct-to-biology approach enables rapid, high-throughput evaluation of large compound libraries in biological assays, eliminating costly and time-consuming purification steps. This strategy has been widely used in the development of proteolysis-targeting chimeras (PROTACs) to facilitate rapid linker optimization and the identification of active degraders from thousands of crude candidate compounds. Similarly, some other direct-to-biology biophysical assays have been utilized for the optimization of small-molecule ligands. However, a direct-to-biology strategy for evaluating cellular target engagement has not yet been demonstrated. Here, we extend this approach to the cellular thermal shift assay (CETSA). By systematically comparing crude reaction mixtures of reported DCAF11 covalent ligands with their corresponding purified analogues, we demonstrate that unpurified compounds can be directly evaluated in CETSA to assess cellular target engagement.

## Introduction

One of the major bottlenecks in the optimization of hit compounds into cell-active chemical probes or drug development candidates is the purification of newly synthesized compounds. In a typical medicinal chemistry campaign, 5–25 mg of each compound is synthesized to ensure sufficient materials for biological evaluation. This requires large quantities of starting materials for synthesis, large volumes of solvents for purification, and entails considerable time for purification, solvent evaporation, and compound handling. In contrast, the direct-to-biology approach enables the parallel synthesis of hundreds to thousands of compounds on the microgram- to nanogram-scale, allowing crude reaction mixtures to be tested directly in biological assays without purification^1^.

Direct-to-biology strategies have been used widely in the development of Proteolysis Targeting Chimeras to identify efficient degraders of target proteins and, to some extent, in other biological assays. Using HiBiT-tagged target proteins, thousands of proteolysis-targeting chimera (PROTAC) candidates have been screened directly in cells with unpurified compounds to rapidly identify active degraders. Subsequent validation with purified samples confirmed activities within one to two orders of magnitude of those observed for crude mixtures, enabling efficient parallel exploration of large linker libraries and accelerating PROTAC discovery^2-5^. Similarly, in affinity selection mass spectrometry (ASMS) assays, nanoscale-synthesized crude compounds produced results consistent with those obtained from the same compounds prepared and purified at the milligram scale^1^. Unpurified compounds have also been successfully employed in other biological assays, such as Microscale thermophoresis (MST) assay^6^, RNA methyltransferase (MTase) assay^7^, surface plasmon resonance (SPR) assay^8^, and fragment hit co-crystallization studies^9^, further demonstrating the versatility of this approach.

However, there have been no reports assessing cellular target engagement using crude samples. Cellular thermal shift assay (CETSA) measures the change in thermal stability of the target proteins upon ligand binding and has been widely used to evaluate the cellular target engagement by the ligands^10-12^. Applying CETSA directly to crude compounds has the potential to accelerate the evaluation of the cellular target engagement of newly synthesized molecules. In the HiBiT-based CETSA assay, the target protein is tagged with the HiBiT peptide. Upon addition of LgBiT in cells, HiBiT complements with LgBiT to form the active NanoLuc luciferase, generating a luminescent signal proportional to the amount of soluble target protein. During thermal challenge, protein unfolding and aggregation lead to loss of soluble protein and a corresponding decrease in luminescence. Ligand binding stabilizes the target protein against thermal denaturation, resulting in increased thermal stability and retention of luminescent signal at elevated temperature This HiBIT-based CETSA assay can be used in plate format to screen a library of compounds which make suitable for direct to biology assay^13^.

Using published DCAF11 covalent ligands as starting points^14^, we synthesized 21 compounds, both crude and with >95% purity, and tested them in the HiBIT-based CETSA assay, demonstrating for the first time that crude compounds can be used in CETSA to distinguish cell-active compounds from inactive ones.

We chose the published DCAF11 covalent ligand, GW5074, as a starting point^14^, as it is commercially available, its derivatives can be made by a simple two-step procedure starting from commercially available reagents, and some structure activity relationship was also reported. PROTAC compounds (e.g., Compound 9, 10,11) based on GW5074 were designed to target BRD4 degradation via the autophagy mechanism, but later were found to be mediated by DCAF11, and no direct cellular engagement of DCAF11 was reported. The structure activity relationship was demonstrated by the ability of GW5074 analogues with ethylene glycol linkers (such as compound 45) to inhibit degradation by the most efficient PROTAC (compound 9). Of all the compounds tested, Compound 45 inhibited the degradation by the PROTAC compound 9 most efficiently (Figure 1) and was used as a reference in this study^14^.

**Figure 1.**
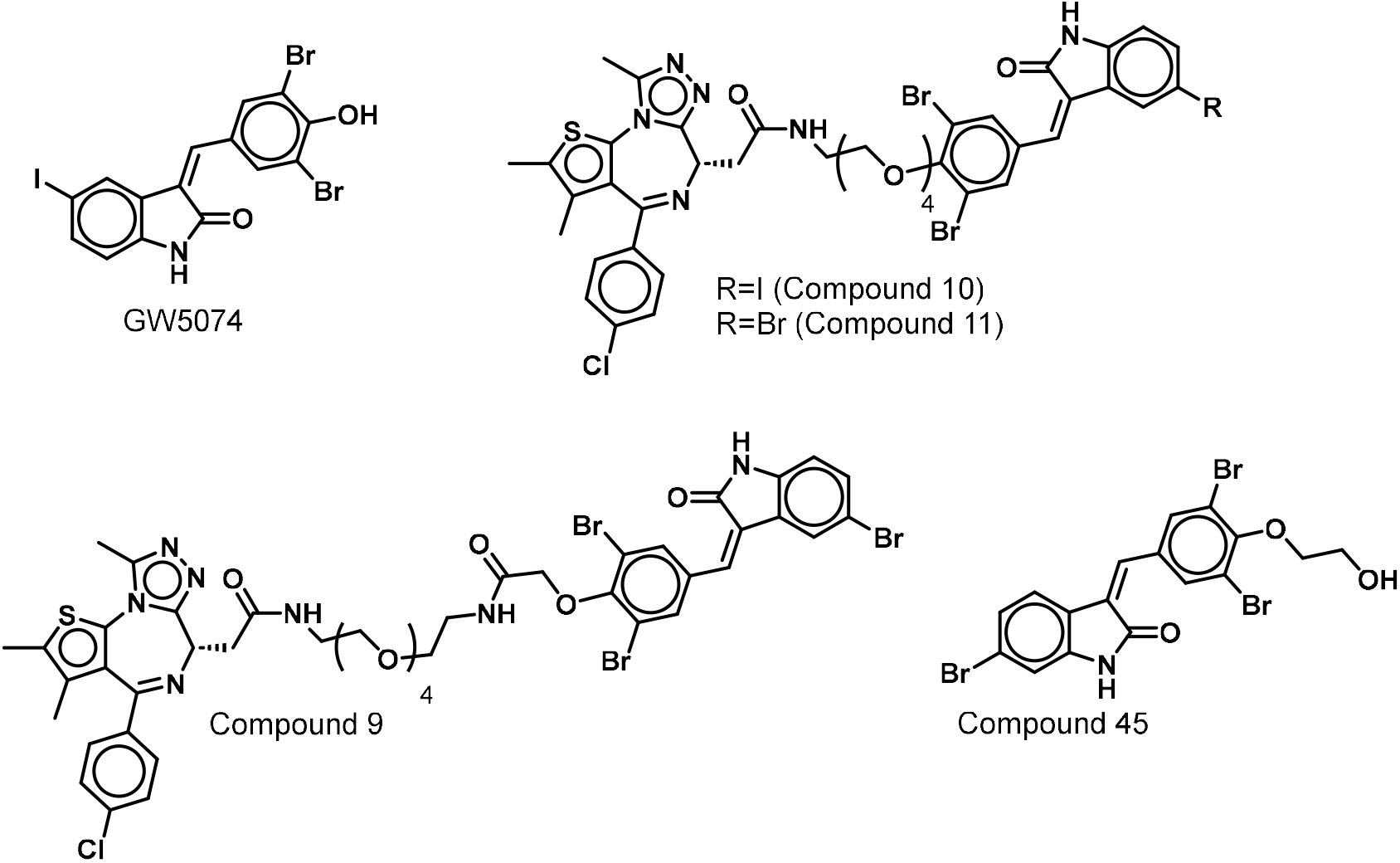
Structures of GW5074, corresponding PROTAC compounds, and an analogue with an ethylene glycol linker

## Results and Discussions

A series of GW5074 analogues with an ethylene glycol linker, including compound 45, was synthesized according to Scheme 1. Substituted 4-hydroxy benzaldehydes (**1a-c**) were alkylated with 2-bromoethanol, and the resulting O-alkylated aldehydes (**2a-c**) were treated with indolinones (**3a-i**) to afford the desired benzylidene indolinones (analogues of GW5074, Table 1) as a mixture of E and Z isomers.

**Scheme 1.**
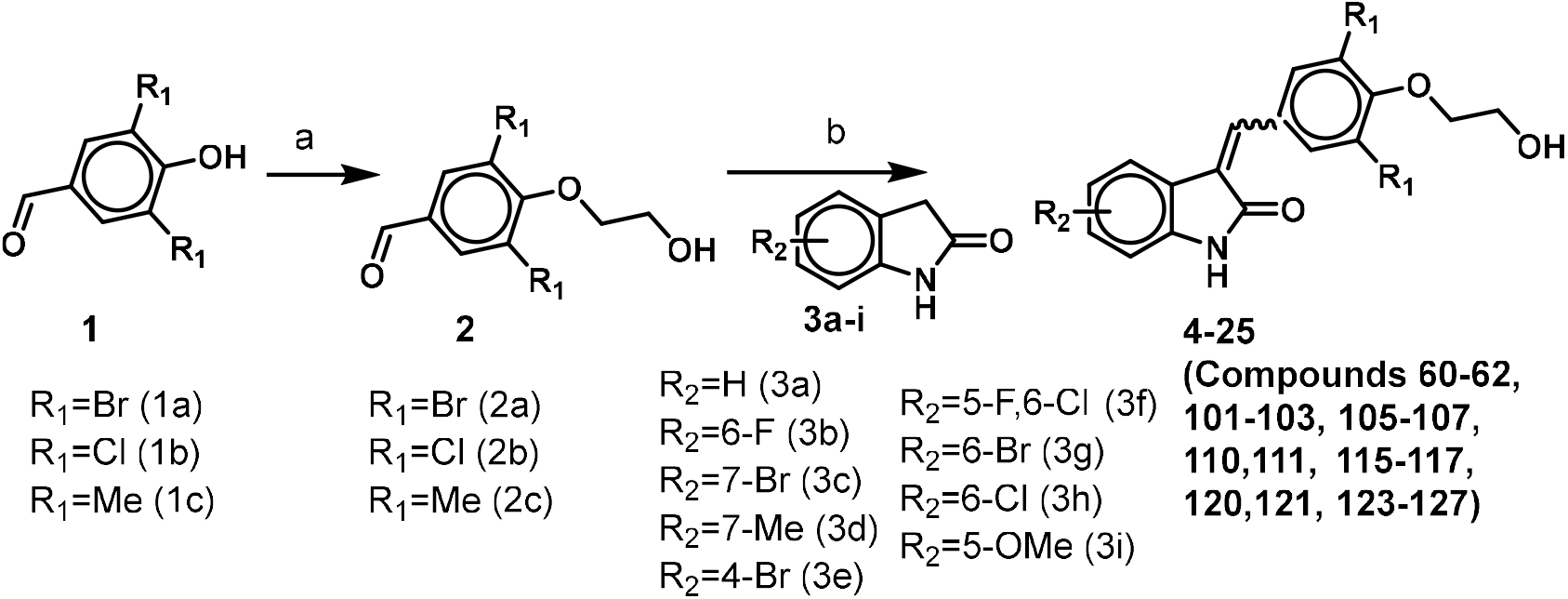
Synthesis of GW5074 analogues. (a) bromoethanol (1.5 eq), K_2_CO_3_ (3 eq), 120° C, 8 h, (b) Piperidine (2 eq) in ethanol, reflux, 12h

**Table 1.** Synthesized and tested analogues of GW5074 and compound 45.

| Compound | $R_1$ | $R_2$ | Compound | $R_1$ | $R_2$ |
| --- | --- | --- | --- | --- | --- |
| 60 (**4**) | Br | H | 115 (**15**) | Cl | 4-Br |
| 61 (**5**) | Cl | H | 116 (**16**) | Cl | 5-F, 6-Cl |
| 62 (**6**) | Me | H | 117 (**17**) | Cl | 6-Br |
| 101 (**7**) | Br | 6-F | 120 (**18**) | Me | 6-Cl |
| 102 (**8**) | Br | 7-Br | 121 (**20**) | Me | 6-F |
| 103 (**9**) | Br | 7-Me | 123 (**21**) | Me | 7-Me |
| 105 (**10**) | Br | 4-Br | 124 (**22**) | Me | 5-OMe |
| 106 (**11**) | Br | 5-F, 6-Cl | 125 (**23**) | Me | 4-Br |
| 107 (Compound 45) (**12**) | Br | 6-Br | 126 (**24**) | Me | 5-F, 6-Cl |
| 110 (**13**) | Cl | 6-Cl | 127 (**25**) | Me | 6-Br |
| 111 (**14**) | Cl | 6-F | GW5074 | Br | 5-I |

We synthesized 21 compounds by varying substitutions on the indolinone and phenyl rings to generate a structurally diverse set of compounds. To assess the compatibility of two commonly used reaction conditions for the synthesis of these compounds in the CETSA assay, the dibromo derivatives were prepared using sodium acetate in acetic acid, and the remaining compounds were synthesized using piperidine in ethanol. In both cases, the reaction mixtures were evaporated to dryness, and the crude products were directly tested in the CETSA assay without further purification or workup, using GW5074 as a reference compound for comparison. In the initial experiment, using GW5074, we found that 57 °C provided the largest assay window in the HiBiT-DCAF11 CETSA assay and was therefore selected for subsequent experiments.

All compounds were tested at 10 µM. Of the 21 compounds tested, 12 compounds, including compound 45 which was the active compound reported^14^, showed significant DCAF11 thermal stabilization relative to DMSO (Figure 2a) and levels comparable to or better than those of the reference compound GW5074. The active compounds were subsequently resynthesized to >95% purity and re-evaluated in the CETSA assay under identical conditions alongside their corresponding crude compounds (Figure 2b). Most of the purified compounds produced results consistent with the corresponding crude mixtures, except compound 125, which exhibited significantly better stabilization than its crude form and showed better stabilization than other pure compounds tested.

**Figure 2.**
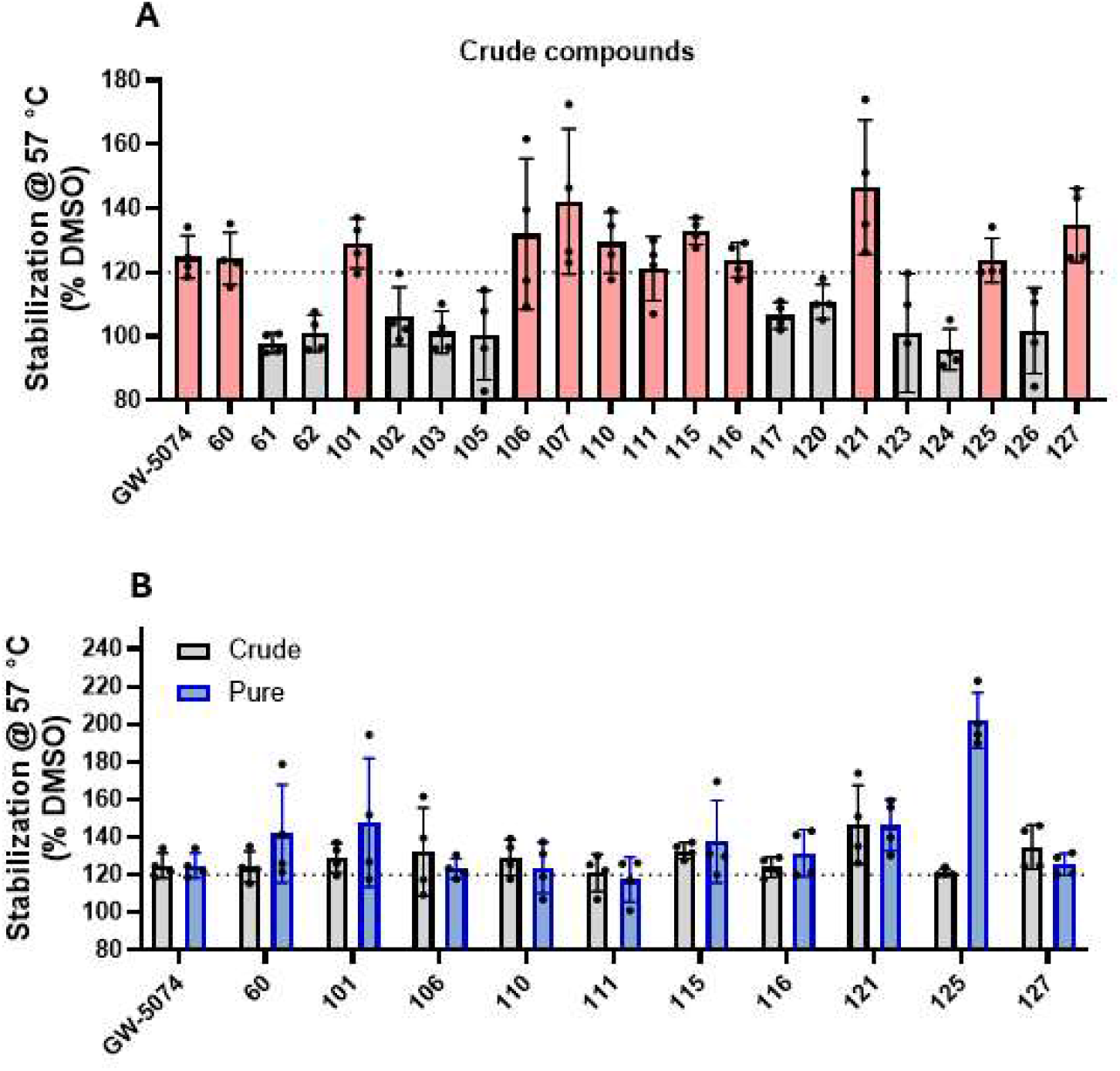
DCAF11 thermal stabilization by analogues of GW5074. HiBIT-DCAF11 overexpressed HEK293T cells were treated with 10 μM compounds for 1 h and heated at 57 ºC for 3 min. (A) DCAF11 stabilization at 57 ºC by crude compounds compared to DMSO. The compounds showing significant stabilization (highlighted in pale red) were selected for resynthesis and purification (B) DCAF11 stabilization by resynthesized compounds (in blue) compared with the corresponding crude compounds (in grey)

To further validate this effect and assess specificity, compound 125 was evaluated at several concentrations, and tested against an unrelated protein, and assessed for cellular toxicity. Compared to DMSO, compound 125 induced dose-dependent thermal stabilization at 10 µM and 20 µM (Figure 3a). At 10 µM, it did not stabilize CBLB, an unrelated protein (Figure 3b), nor did it affect cell growth in three different cell lines (Figure 3c), confirming the specificity of the observed effect.

**Figure 3.**
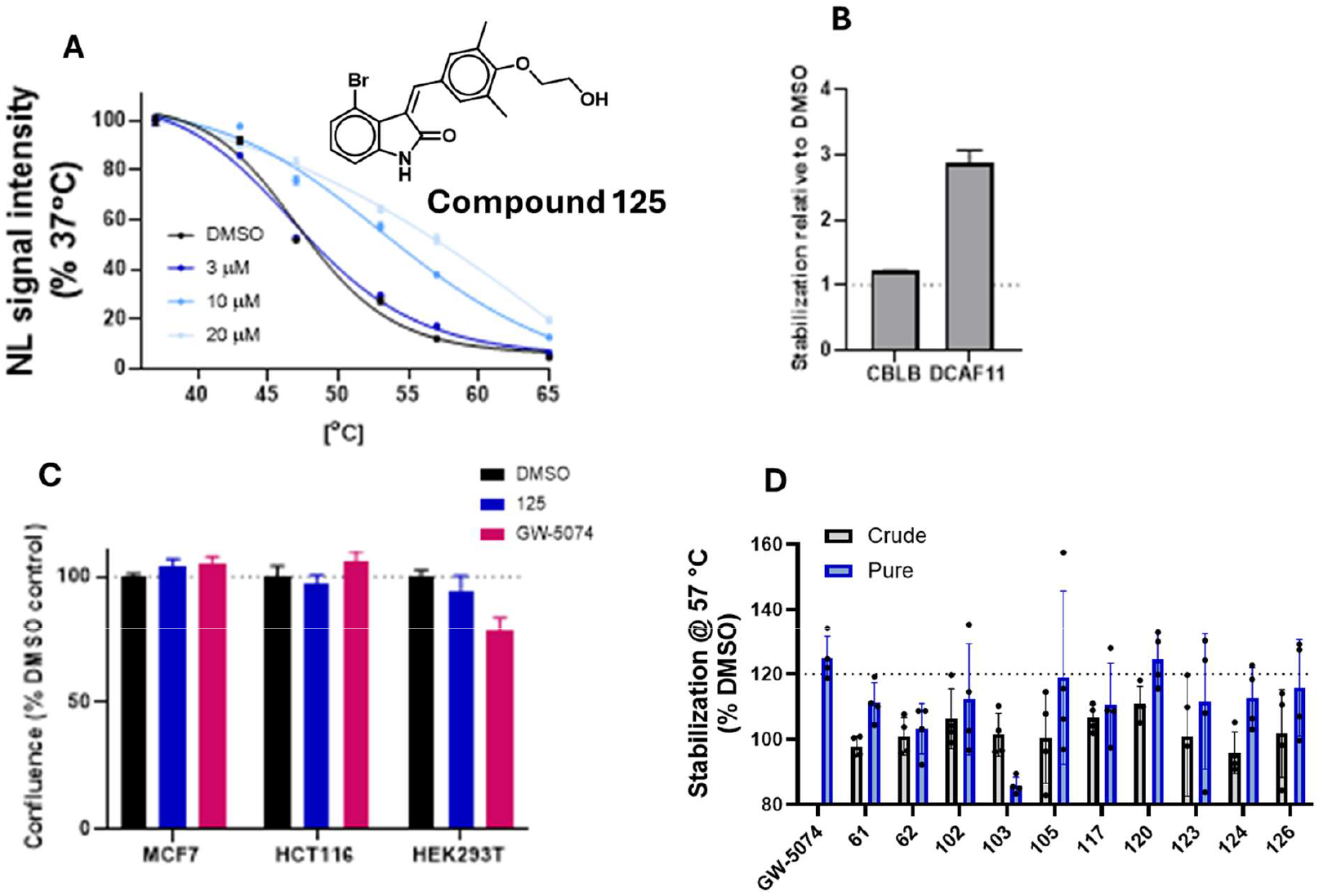
Compound 125 selectively stabilizes DCAF11 in HEK293T cells. Cells were treated with compounds for 1 h and heated at the indicated temperatures for 3 min. (A) Dose response of compound 125 by HiBIT CETSA (B), Compound 125 (10 μM) treatment increases stability of DCAF11 but not unrelated protein CBLB (C), The effect of compounds treatment (10 μM) on cell growth (84 h). (D), Thermal stabilization profiles of analogues of GW5074 (10 μM),which crude mixtures did not show significant thermal stabilization of DCAF11, resynthesized compounds (in blue), crude compounds (in grey).

Because purified compound 125 exhibited greater thermal stabilization than the crude sample, we sought to ensure that no potentially active compounds had been overlooked when tested as crude mixtures. Following the same approach applied to the active compounds, all previously inactive compounds were resynthesized and purified, and then evaluated in the CETSA assay alongside their corresponding crude samples. None of the purified compounds demonstrated greater stabilization than the reference compound GW5074, confirming the same trend observed with crude compounds (Figure 3d). Although there are some structure activity relationship trends, the CETSA assay - especially when used in a direct-to-biology format - cannot be used for determining quantitative structure activity relationships, and is limited to assessing cellular target engagement. While completing this study, a more potent PROTAC compound with the same benzylidene indolinone was reported and not included in this direct to biology studies^15^.

## Conclusion

By comparing DCAF11 protein stabilization levels induced by crude samples of the analogues of the reported DCAF11 ligand GW5074 with those obtained from the corresponding purified samples, we demonstrated that cellular target engagement can be reliably assessed using crude compounds alone in the HiBIT-based CETSA assay. Applying this strategy, we identified a compound (compound 125) with improved cellular target engagement relative to GW5074. This study highlights the broader applicability of the CETSA assay for evaluating target engagement within a direct-to-biology framework, enabling rapid assessment of cellular activity and thereby accelerating the development of cell-active chemical tools.

## Methods

### COMPOUND SYNTHESIS

#### General Conditions

Analytical HPLC analyses were carried out on Waters Aquity series instrument. A linear gradient starting from 5% acetonitrile and 95% water (0.1% formic acid) to 95% acetonitrile and 5% water (0.1% formic acid) over 2 minutes, followed by 1 minutes of elution at 95% acetonitrile and 5% water (0.1% formic acid) was employed. Flow rate was 1 mL/min, and UV detection was set to 254 nm and 214 nm. All compounds submitted for testing were at ≥95% purity (by HPLC, by UV detection at 254nm) unless otherwise stated. Purities of all compounds were estimated to be >95%.

#### Intermediates 2a-C

To a solution of Hydroxybenzaldehyde (5 mmol, 1 eq.) and bromoethanol (7.5 mmol, 1.5 eq.) in DMF (10 mL) was added K_2_CO_3_ (3 eq, 15 mmol). The mixture was heated under microwave conditions at 120° C for 8 h. Crude material was transferred into a separating funnel with water (40 ml). The reaction mixture was extracted with EtOAc (2 × 20 mL). The organic layers were combined, washed with brine (2 × 10 mL), dried over anhydrous MgSO_4_, filtered, and concentrated. The crude product was purified by silica gel column chromatography with hexanes/EtOAc.

#### Compounds 60, 101-103, 105-107

A solution of 3’,5’-dibromo-4’-hydroxybenzaldehyde (1 mmol, 1 eq.), the indolinones (1.1 mmol), and anhydrous sodium acetate (2 mmol) in glacial acetic acid (10 mL) was refluxed for 12 h. Upon completion, the reaction mixture was cooled to room temperature and concentrated under reduced pressure, which was used without further purification in the direct to biology assay. The resulting residue was poured onto crushed ice, and the solid that formed was collected by filtration. The crude product was purified by recrystallization from methanol.

#### Compounds 61,62, 110,111, 115-117, 120-121, 122-127

3’,5’-dichloro or dimethyl-4’-hydroxybenzaldehyde (1 mmol, 1 eq.) was dissolved in ethanol (10 mL), followed by the addition of the corresponding indolinones (1.1 mmol) and piperidine (2 mmol, 2 eq.). The mixture was refluxed for 12 h and then cooled to room temperature and concentrated under reduced pressure, which was used without further purification in the direct to biology assay. The crude products were purified by recrystallization from methanol.

### HiBIT CETSA

The pBiT3.1-C vector (Promega) was used to generate C-terminally HiBiT-tagged DCAF11 or CBLB constructs. HEK293T cells were seeded at 2 × 10^5^ cells per 6-well plate in DMEM supplemented with 10% FBS, 100 U/mL penicillin, and 100 μg/mL streptomycin. Cells were transfected with 0.2 μg of target plasmid and 1.8 μg of empty vector using X-tremeGENE™ HP DNA Transfection Reagent (Roche) according to the manufacturer’s instructions and incubated for 24 h.The following day, cells were trypsinized, centrifuged at 300 × g for 2 min, and resuspended in Opti-MEM™ I Reduced Serum Medium (no phenol red) at a final density of 2 × 10^5^ cells/mL. Cells were transferred to a 96-well PCR plate (40 μL per well; one well per temperature condition), treated with compounds or DMSO control, and incubated for 1 h at 37 °C and 5% CO_2_. Thermal challenge was performed in a gradient thermocycler using the following program: 1 min at 22 °C, 3 min at the indicated temperature, and 2 min at 22 °C. Subsequently, 40 μL per well of lysis buffer (2% NP-40, protease inhibitors, Opti-MEM™ I Reduced Serum Medium without phenol red) containing 200 μM LgBiT was added, and samples were incubated for 10 min at room temperature. After addition of 20 μL per well of 100-fold diluted NanoBRET™ Nano-Glo® substrate (Promega) prepared in Opti-MEM™ I Reduced Serum Medium (no phenol red), samples were mixed, and 20 μL per well was transferred into 384-well white plates in technical quadruplicates using a multichannel pipette. Bioluminescence was measured using a CLARIOstar plate reader.

### Cell growth assay

To assess the effect of compounds on cell growth, HEK293T, MCF7, and HCT116 cells were seeded in 96-well plates and treated with compounds at a final concentration of 10 µM. Live-cell imaging was performed using an Incucyte live-cell analysis system (Sartorius) over a period of 0–84 h. Cell confluence was quantified using the Incucyte Software (2023A V2).

## ACKNOWLEDGEMENTS

This research was undertaken thanks in part to funding provided to the University of Toronto’s Acceleration Consortium from the Canada First Research Excellence Fund. Grant number - CFREF-2022-00042. The Structural Genomics Consortium is a registered charity (no: 1097737) that receives funds from Bayer AG, Boehringer Ingelheim, Bristol Myers Squibb, Genentech, Genome Canada through Ontario Genomics Institute [OGI-196], EU/EFPIA/OICR/McGill/KTH/Diamond Innovative Medicines Initiative 2 Joint Undertaking [EUbOPEN grant 875510], Janssen, Merck KGaA (aka EMD in Canada and US), Pfizer and Takeda.

## Author Contributions

The manuscript was written through contributions of all authors. All authors have given approval to the final version of the manuscript. DP synthesized, and MS and SP tested the compounds, AI generated the cell lines. Guidance and advice throughout the project were provided by DB, RB and VS.

